# Interrogation of dynamic glucose-enhanced MRI and fluorescence-based imaging reveals a perturbed glymphatic network in Huntington’s disease

**DOI:** 10.1101/2023.04.03.535397

**Authors:** Hongshuai Liu, Lin Chen, Chuangchuang Zhang, Chang Liu, Yuguo Li, Liam Cheng, Zhiliang Wei, Ziqin Zhang, Hanzhang Lu, Peter C. M. van Zijl, Jeffrey J. Iliff, Jiadi Xu, Wenzhen Duan

## Abstract

Huntington’s disease (HD) is a neurodegenerative disorder that presents with progressive motor, mental, and cognitive impairment leading to early disability and mortality. The accumulation of mutant huntingtin protein aggregates in neurons is a pathological hallmark of HD. The glymphatic system, a brain-wide perivascular network, facilitates the exchange of interstitial fluid (ISF) and cerebrospinal fluid (CSF), supporting interstitial solute clearance including abnormal proteins from mammalian brains. In this study, we employed dynamic glucose-enhanced (DGE) MRI to measure D-glucose clearance from CSF as a tool to assess CSF clearance capacity to predict glymphatic function in a mouse model of HD. Our results demonstrate significantly diminished CSF clearance efficiency in premanifest zQ175 HD mice. The impairment of CSF clearance of D-glucose, measured by DGE MRI, worsened with disease progression. These DGE MRI findings in compromised glymphatic function in HD mice were further confirmed with fluorescence-based imaging of glymphatic CSF tracer influx, suggesting an impaired glymphatic function in premanifest stage of HD. Moreover, expression of the astroglial water channel aquaporin-4 (AQP4) in the perivascular compartment, a key mediator of glymphatic function, was significantly diminished in both HD mouse brain as well as postmortem human HD brain. Our data, acquired using a clinically translatable MRI approach, indicate a perturbed glymphatic network in the HD brain as early as in the premanifest stage. Further validation of these findings in clinical studies should provide insights into potential of glymphatic clearance as a HD biomarker and for glymphatic functioning as a disease-modifying therapeutic target for HD.

## INTRODUCTION

Huntington’s disease (HD) is a dominantly inherited, fatal neurodegenerative disorder caused by a CAG expansion in the exon-1 of the *Huntingtin* (*HTT*) gene which leads to the production of a toxic mutant huntingtin protein (mHTT) (*1*). A fundamental pathological hallmark of HD is abnormal mHTT accumulation in the brain, where the misfolded and aggregated disease-causative protein propagates and spreads in a prion-like fashion (*2*).

A tenet of brain homeostasis is that protein clearance must equal protein synthesis. Interstitial solutes, including proteins, can be cleared across the blood-brain barrier and undergo local cellular uptake and degradation. When not subject to these cellular processes, it was generally believed that they were cleared through exchange with the surrounding cerebrospinal fluid (CSF) in a process that was considered slow and diffuse. Beginning in 2012, the glymphatic (glial-lymphatic) model of fluid and solute exchange was described (*3*), in which CSF and interstitial fluid (ISF) and solute exchange were observed to be rapid, anatomically organized along perivascular spaces surrounding the cerebral vasculature, and regulated by the sleep-wake cycle (*4*). More recently, meningeal lymphatic vessels associated with dural sinuses have been characterized that contribute to the clearance of solutes, including those arising from the brain interstitium, from the CSF compartment (*5-7*). The combined activity of perivascular glymphatic exchange and meningeal lymphatic clearance appear to function together to subserve brain interstitial solute and waste clearance. Importantly, key features of these processes, including perivascular CSF-ISF solute exchange, sleep-active brain solute clearance, and parasagittal solute uptake have been confirmed in the human brain (*8*).

The glymphatic system is believed to be comprised of meningeal lymphatic vessels that drain CSF and ISF toward cervical lymph nodes(*4*). The function of the glymphatic system is supported by aquaporin-4 (AQP4) water channel, which presents with high density in perivascular astrocytic endfoot membranes (*4*). Glymphatic function is a highly regulated process with changes in its activity accompanying aging as well as disease conditions (*9-11*). The efficiency of glymphatic clearance is lowered when AQP4 perivascular localization is reduced (*12, 13*). There is a growing awareness that such reduced AQP4 perivascular localization occurs in neurological disorders (*14-17*), which in turn associated with the expression of a protein complex including α-syntrophin (SNTA1) (*18, 19*).

Evidence has begun to emerge that failure of the glymphatic system leads to an increase of local mHTT concentrations to levels that favor aggregation. One recent study suggested that a dysfunctional glymphatic system may contribute to HD manifestation and interrupt antisense oligonucleotide (ASO) distribution throughout the entire brain (*20*). Another study reports that secretion of mHTT from cells in the brain, followed by glymphatic clearance from the extracellular space, contributes to mHTT in the CSF (*21*). Whether the glymphatic system is disturbed in the HD brain, particularly in the premanifest stage, remains unclear.

In this study, we combined an *in vivo* measure of CSF clearance capacity by dynamic glucose-enhanced (DGE) MRI (*22*) with fluorescence-based imaging of glymphatic CSF tracer influx in a widely-used zQ175 knock-in HD mouse model. We observed that CSF clearance efficiency and glymphatic function were impaired in the zQ175 HD mice, prior to the manifestation of motor deficits and detectable striatal atrophy. The impairment of CSF clearance worsened along HD progression. Further mechanistic study indicated that AQP4 perivascular localization, a key contributor to glymphatic function, was significantly reduced in both HD mice and HD human brain. These findings set the premise to further investigate the role of glymphatic function in HD pathogenesis as a potential therapeutic target as well as an early biomarker for this devastating disease.

## RESULTS

### CSF clearance capacity measured by DGE MRI is AQP4-dependent and reflects glymphatic CSF-ISF exchange capacity

We recently developed a DGE MRI (*22-25*) approach that can sensitively assess D-glucose signal changes in the mouse CSF (*26*). Because glucose transporters are highly enriched in the brain capillary endothelial cells and choroid plexus epithelial cells (*27, 28*), D-glucose can rapidly penetrate the brain CSF barrier (BCSFB) and enter CSF. This unique feature provides an opportunity to measure CSF clearance efficiency through monitoring D-glucose levels in CSF following intravenous delivery. Because AQP4 (protein) is essential for functional glymphatic transport (*29*) and significantly compromised glymphatic function has been reported in *Aqp4* KO mice (*30*), we first evaluated CSF clearance efficiency in an *Aqp4* knock-out mouse model.

We confirmed levels of AQP4 in the KO mouse brains (**Figure S1A**), then conducted DGE MRI scans in both 4-month-old *Aqp4* KO mice and the wild type (WT) mice. Following D-glucose injection via the tail vein, DGE signal in the CSF rapidly reached a peak in both mouse genotypes. In WT mice, D-glucose signal decayed gradually following the peak, indicating gradual D-glucose clearance from CSF, while DGE signal in the CSF of *Aqp4* KO mice was sustained or even slightly elevated during the MRI scan session (40min) (**Figures S1B, S1C**). The uptake of D-glucose in CSF was comparable (p>0.05) in the two groups (**Figure S1D**), while D-glucose clearance was significantly lower (p < 0.05) in Aqp4 KO mice than WT mice (**Figure S1E**). Here we used a γ-variate model with three unknowns (S_max_, μ_in_, and μ_out_) to fit the uptake and get an initial rate estimate (μ_out_) for the clearance by fitting only the first 20-min post-infusion. This method from our previous studies(*26*) is summarized in the supplement. Taken together, these results suggest that loss of the *Aqp4* gene, which suppresses perivascular glymphatic exchange, slows CSF D-glucose clearance. Thus, CSF D-glucose clearance capacity is AQP4 dependent.

In subsequent experiments (detailed below), we evaluated whether altered CSF clearance efficiency paralleled changes in the influx of fluorescent CSF tracers into brain tissue of HD mice, a validated approach for assessing glymphatic function in rodents. Importantly, similar results were observed. Combined, these results strongly support the notion that CSF D-glucose clearance efficiency reflects glymphatic CSF-ISF exchange capacity.

### An impaired glymphatic function is evident in premanifest zQ175 HD mice from combined measures of (i) CSF clearance efficiency by DGE MRI and (ii) influx of a fluorescence CSF tracer from cisterna magna to brain parenchyma

Using structural MRI measures and motor function assessment, we confirmed that there was no significant striatal atrophy (**Figure S2A**), and motor function on the balance beam (**Figure S2B**) was preserved in 4-month-old zQ175 (heterozygous) HD mice, indicating that this age is a premanifest stage of zQ175 HD model.

We then used DGE MRI to assess CSF D-glucose clearance in premanifest zQ175 HD mice. After a baseline DGE signal was established, D-glucose was injected via the tail vein into 4-month-old zQ175 mice and WT littermate controls, and DGE MRI scans were conducted over 40 min period. We monitored DGE signals in the CSF of the third ventricle (representative images shown in **Figure 1A**). DGE signals reached a peak in the CSF within 1-2 min following i.v injection of D-glucose. WT mice showed a gradual DGE signal decay, indicating normal D-glucose clearance in the CSF. In contrast, zQ175 HD mice exhibited sustained D-glucose signals in the CSF for the entire scan period (**Figures 1A, 1B**) (*p*<0.05 vs. WT). The D-glucose clearance in zQ175 HD mice was significantly slower than that in WT mice (p<0.05, **Figure 1C**), while the D-glucose uptake in CSF was comparable between groups (p>0.05, **Figure 1D**). These CSF D-glucose uptake results are consistent with the levels of glucose transporters in the brain (**Figure S2C**).

**Figure 1.**
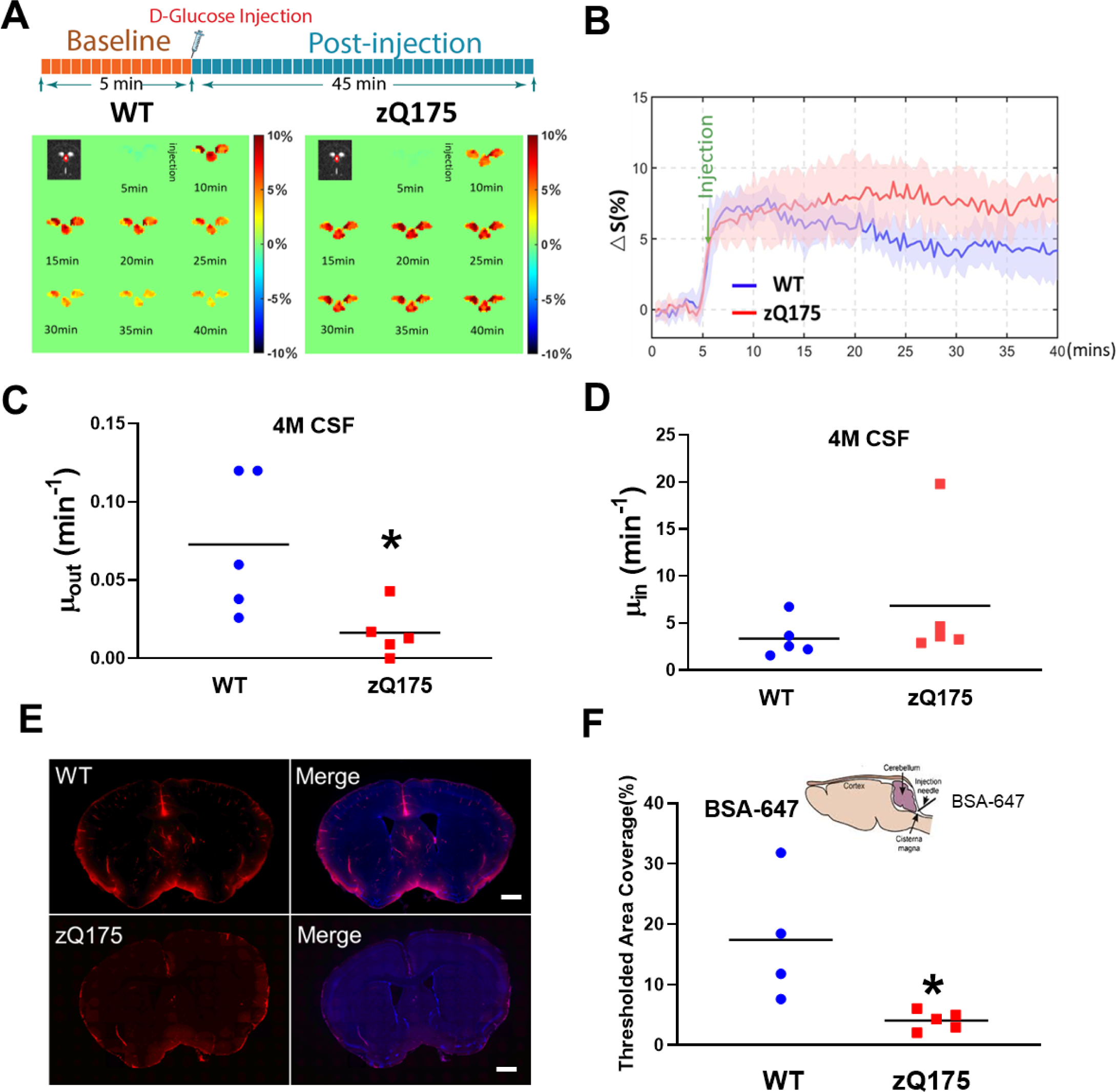
Impaired glymphatic system as revealed by DGE MRI and fluorescence-based imaging in premanifest zQ175 HD mice. (**A**) Illustration of DGE MRI scan timeline (upper panel) and representative DGE images for the 3^rd^ ventricle (lower panel) in a wild type (WT) mouse and a zQ175 mouse at 4 months of age. (**B**) Average CSF-based DGE signal changes during the entire scan period from WT and zQ175 mice. n=5 mice/genotype. (**C**) Comparison of fitted clearance parameter μout for CSF. \**p*<0.05 vs. WT by Standard Student’s t-test. (**D**) Comparison of fitted uptake parameter μin for CSF. (**E**) Representative images of BSA-647 fluorescent dye distribution in the brain parenchyma at 60 min after intra-cisterna magna injection. Note the wide distribution of fluorescent dye along the glymphatic pathway in WT mice, while very limited fluorescent distribution was seen in the HD mouse brain. Scale bar =500 µm. (**F**) Quantification of the fluorescent dye distribution in CSF. \**p*<0.05 vs. WT by Standard Student’s t-test.

To determine whether blood D-glucose levels influenced D-glucose uptake/clearance in the CSF, dynamic glucose signals in the sagittal sinus were monitored during the entire scan period. We observed no significant difference in blood DGE signals between the two genotypes (**Figure S2E**), including glucose clearance, uptake, and maximal D-glucose signal difference in the vein (**Figures S2F-H**). These results indicate that the CSF data from Figure 1 suggest a potential impairment of glymphatic function in premanifest zQ175 HD mice.

We further validated these MRI findings with a gold-standard fluorescence-based CSF tracer imaging technique. A fluorescent tracer Alexa Fluor 647 labeled BSA (BSA-647) was injected into the cisterna magna in premanifest zQ175 mice; sixty minutes following tracer injection, BSA-647 fluorescent dye distribution in the parenchyma was quantified. We observed significantly reduced fluorescent tracer movement into the brain parenchyma along the glymphatic network in 5-month-old zQ175 HD mice compared to age-matched WT controls (**Figures 1E, F**), confirming the impaired glymphatic function in premanifest zQ175 HD mice. Taken together, these data show the capacity of DGE MRI to detect glymphatic function and that glymphatic function is compromised in zQ175 HD mice prior to striatal atrophy and motor deficits.

### Reduced AQP4 perivascular localization in premanifest zQ175 HD mouse brain

Glymphatic clearance efficiency is mediated by astrocytic AQP4 water channels that are present at high density in the perivascular astrocytic endfoot membranes. Thus, AQP4-dependent convective flow is critical for effective glymphatic clearance, and AQP4 perivascular localization is required for glymphatic function. To determine if AQP4 perivascular localization was altered in the 4-month-old zQ175 HD mouse brain, we performed co-immunostaining of AQP4 and vascular marker protein collagen IV to examine AQP4 perivascular colocalization. As visualized in **Figure 2**, AQP4 immunosignals were concentrated on the perivascular domains in both the striatum (**Figure 2A**) and cerebral cortex (**Figure 2C**) of WT mice, indicated by colocalized AQP4 and collagen IV immunosignals (yellow). In contrast, zQ175 HD mice exhibited significantly reduced AQP4 immunosignals in the perivascular domains in both the striatum (**Figures 2A, B**) and cerebral cortex (**Figures 2C, D**), suggesting loss of AQP4 perivascular localization in the premanifest zQ175 HD mouse brain.

**Figure 2.**
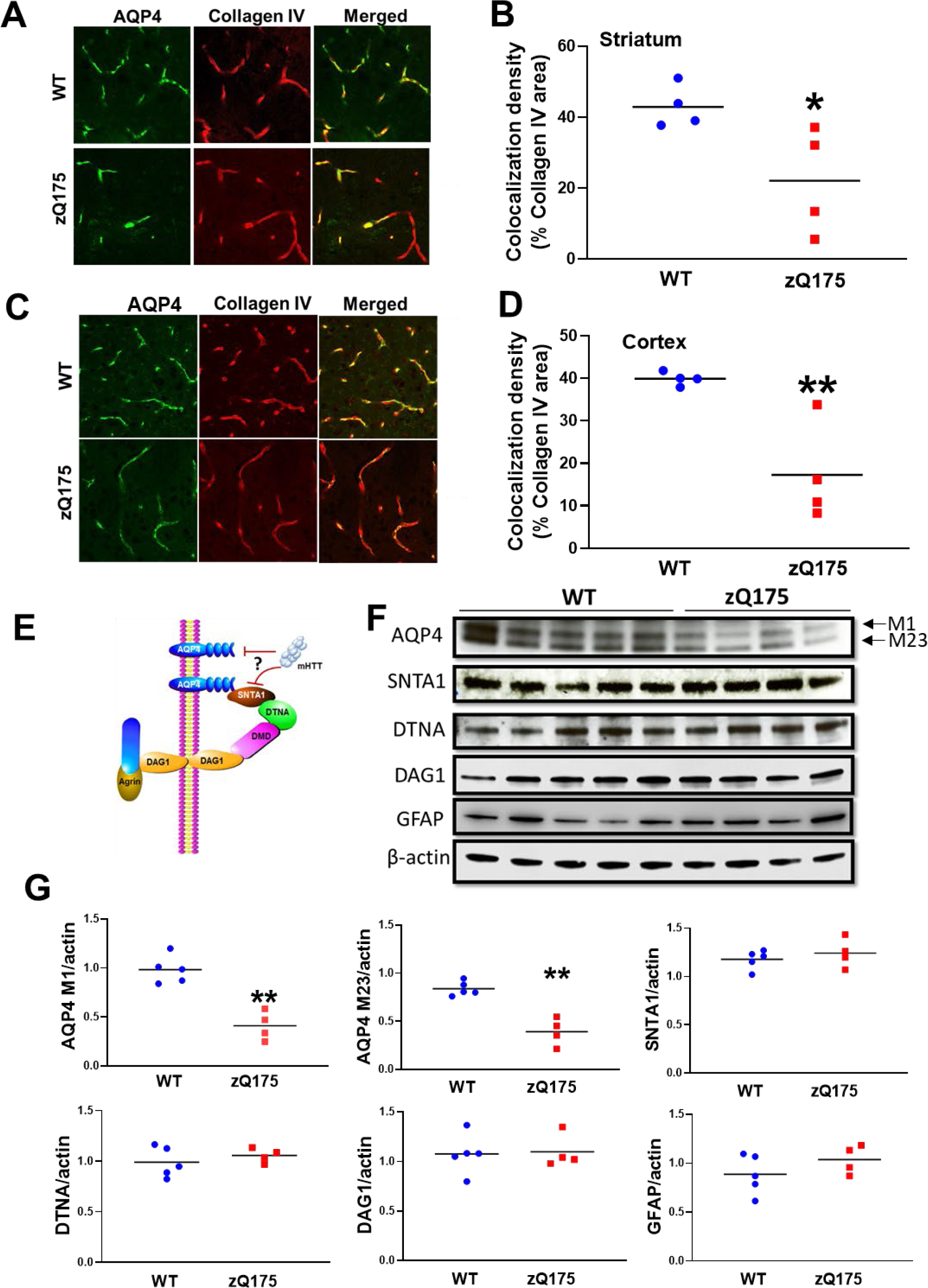
Decreased AQP4 perivascular localization (polarization) and levels in the brain of premanifest zQ175 HD mice. (**A**) Representative images of co-immunofluorescent staining of AQP4 and collagen IV in the striatum of 4-month-old zQ175 and WT controls. (**B**) Quantification of co-localized pixels of AQP4 (green) and collagen IV (red) in the striatum of 4-month-old zQ175 mice and WT controls. \**p*<0.05 vs. WT by Standard Student’s t-test. (**C**) Representative images of co-immunofluorescent staining of AQP4 and collagen IV in the cortex of 4-month-old zQ175 and WT controls. (**D**) Quantification of co-localized pixels of AQP4 (green) and collagen IV (red) in the cortex of 4-month-old zQ175 mice and WT controls. \**p*<0.05 vs. WT by Standard Student’s t-test. (**E**) Illustration of AQP4 and its astrocytic endfeet anchoring protein complex. (**F**) Western blots of AQP4, AQP4 anchoring protein complex components SNTA1, DTNA, DAG1, activated astrocyte marker GFAP, and loading control β-actin. (**G**). Quantification of Western blotting results by Imaging J. \*\**p*<0.01 vs. WT by Standard Student’s t-test.

AQP4 polarization requires the presence of a functional protein complex formed by a key protein SNTA1, which connects AQP4 with other complex proteins, dystrobrevin alpha (DTNA) and dystroglycan (DAG1) (*30*) (**Figure 2E**), and this protein complex anchors AQP4 to the astrocytic endfeet perivascular location. The absence of the key component protein SNTA1 interferes with AQP4 perivascular localization,(*18*) which may disrupt glymphatic function. To determine whether loss of AQP4 perivascular localization was due to reduced levels of AQP4 or its astrocytic endfeet anchoring machinery, we examined protein levels of AQP4, SNTA1, DTNA, and DAG1 in the striatum of premanifest zQ175 mice. There are two major isoforms of AQP4 in the mouse brain, M1 and M23. We found that both isoforms of AQP4 protein were significantly (p<0.01) reduced in the premanifest zQ175 mouse striatum (**Figure 2F, G**). There was no significant difference in the protein levels of all three AQP4 anchoring proteins at this age (**Figure 2F, G**). Because AQP4 is preferentially expressed in astrocytes in the brain, we also examined GFAP levels, and our results indicated no detectable changes in GFAP levels (**Figure 2F, G**). These data suggest that the loss of AQP4 perivascular localization in premanifest zQ175 HD mice is mainly due to reduced AQP4 protein levels.

### Perturbed glymphatic function and further reduced AQP4 perivascular localization in manifest zQ175 HD mice

Like HD patients, zQ175 HD mice progressively develop into the symptomatic (manifest) stage, displaying motor deficits and striatal atrophy (*31, 32*). Using structural MRI measures and motor testing on a balance beam, we confirmed that 10-month-old zQ175 HD mice exhibited significant (p<0.05, t-test) striatal atrophy (**Figure S3A**) and motor deficits, indicated by prolonged traverse time (p<0.05) on the beam compared to their age-matched WT control mice (**Figure S3B**). We then sought to determine whether glymphatic dysfunction progresses along with disease progression in zQ175 mice. We conducted DGE MRI scans in 10-month-old zQ175 mice. zQ175 mice exhibited slower DGE signal decay in the CSF during an entire scan period, while WT mice still performed well in the CSF clearance of D-glucose (**Figures 3A, B**). Further imaging analysis indicated that CSF D-glucose clearance was significantly slower (p<0.01) in the zQ175 than in WT (**Figure 3C**), while CSF D-glucose uptake had no significant difference between the two genotypes (**Figure 3D**). These data suggest that manifest zQ175 HD mice have exacerbated the disruption of glymphatic function.

**Figure 3.**
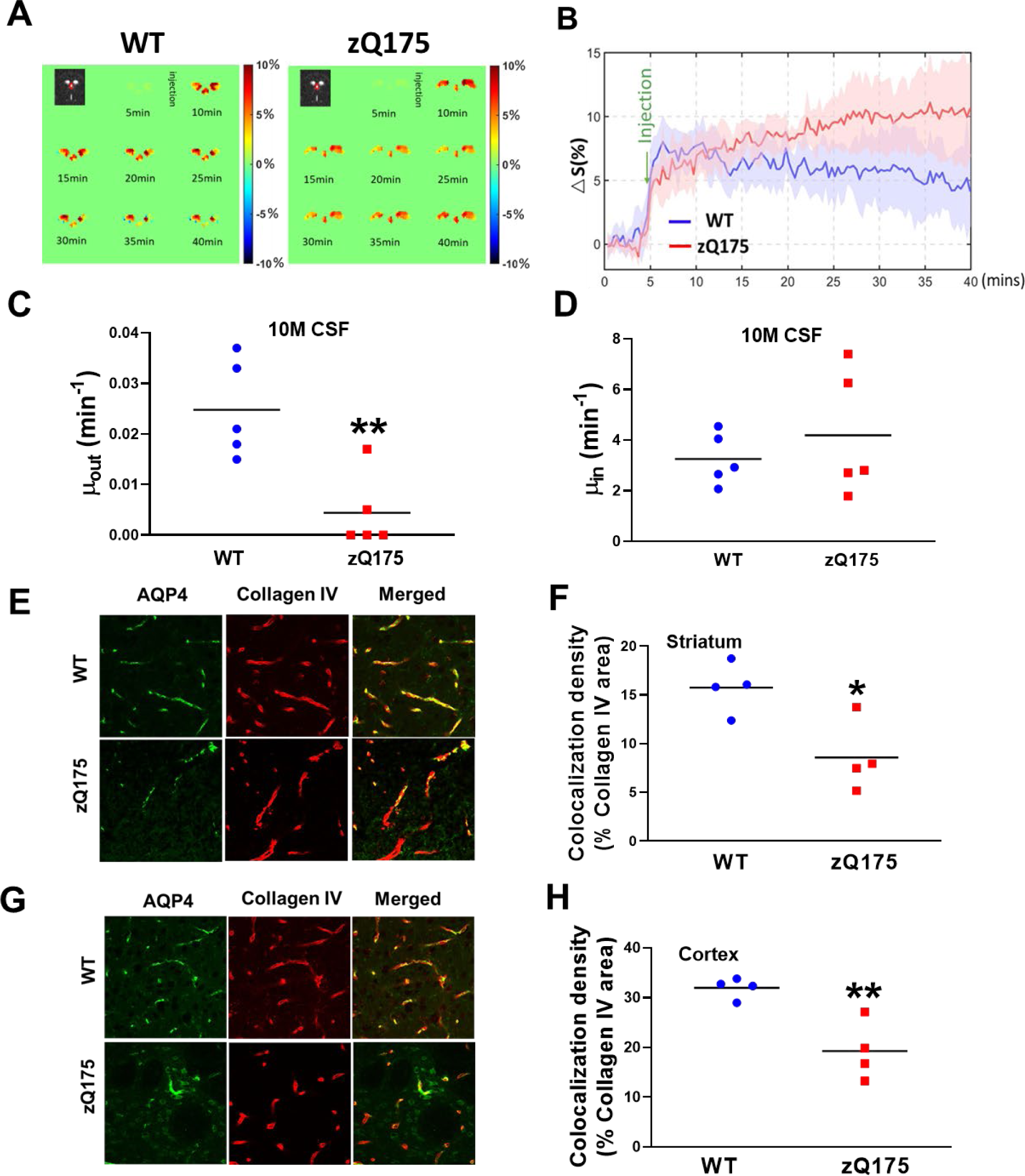
Perturbed glymphatic function and exacerbated AQP4 perivascular localization (polarization) in the manifest zQ175 HD mouse brain. (**A**) Representative images of DGE signals (lower panel) in a wild type (WT) mouse and a zQ175 mouse at 10 months of age. (**B**) Average dynamic D-glucose signals during the entire scan period from WT and zQ75 mice. n=5 mice/genotype. (**C**) Comparison of fitted clearance parameter μout D-glucose clearance rate in CSF of the 3rd ventricle \*\**p*<0.01 vs. WT by Standard Student’s t-test. (**D**) Comparison of fitted clearance parameter μin D-glucose uptake rate in CSF of the 3rd ventricle (**E**) Representative images of co-immunofluorescent staining of AQP4 and collagen IV in the striatum of 10-month-old zQ175 and WT controls. (**F**) Quantification of co-localized pixels of AQP4 (green) and collagen IV (red) in the striatum of 10-month-old zQ175 mice and WT controls. \**p*<0.05 vs. WT by Standard Student’s t-test. (**G**) Representative images of co-immunofluorescent staining of AQP4 and collagen IV in the cortex of 10-month-old zQ175 and WT controls. (**H**) Quantification of co-localized pixels of AQP4 (green) and collagen IV (red) in the cortex of 10-month-old zQ175 mice and WT controls. \*\**p*<0.01 vs. WT by Standard Student’s t-test.

We then asked whether D-glucose uptake in the parenchyma of manifest zQ175 HD mice was altered. Western blotting analysis of glucose transporter 1 (Glut1) and 3 (Glut3) in the striatum indicated that there were no significant changes in levels of Glut1, which is a major glucose transporter expressed in glia and endothelial/epithelial cells (**Figures S3C, D**). Levels of Glut3, which is preferentially expressed in the neurons, were decreased in the manifest zQ175 mouse striatum (p<0.01, **Figures S3C, E**). Consistent with Western blot findings, we observed significantly lower D-glucose uptake in the striatum of manifest zQ175 mice compared to age-matched controls (p<0.05, **Figures S3 F-H**). Taken together, reduced D-glucose uptake in the zQ175 HD mouse striatum is likely due to reduced neuronal Glut3 levels. No changes in Glut1 protein levels are consistent with no changes in D-glucose uptake in the CSF.

Given the critical role astrocytic AQP4 perivascular localization plays in the control of glymphatic function, we performed co-immunostaining with antibodies to AQP4 and collagen IV. Supporting our imaging findings on impaired glymphatic function detected by DGE MRI, AQP4 perivascular localization in the manifest zQ175 mouse brain was significantly diminished, indicated by decreased colocalization of AQP4 and collagen IV in the striatum (p<0.05, **Figures 3E, F**) and cortex (p<0.01, **Figures 3G, H**).

To dissect the underlying molecular basis of reduced AQP4 perivascular in manifest zQ175 HD mice, we quantified the levels of AQP4 and its astrocytic endfeet anchoring protein SNTA1 in the mouse striatum at 10 months of age. When we quantified levels of these proteins using a ratio to β-actin, we found that AQP4 M1, M23, and SNTA1 levels had no significant difference between HD mice and age-matched controls (**Figures S4A, B**). Meanwhile, we noticed increased GFAP levels in the zQ175 HD mice at the manifest age (**Figures S4A, B**), indicating astrogliosis in the HD mouse brain. Because AQP4 and SNTA1 are exclusively expressed in the astrocytes, we asked whether AQP4 and SNTA1 levels are proportionally altered in astrocytes. By calculating the ratio of these proteins to GFAP, we observed that the ratio of both isoforms of AQP4, M1 and M23, to GFAP was decreased in manifest zQ175 mice (p<0.05, **Figure S4C**). Notably, SNTA1 levels had a trend to decrease but were not statistically different (**Figure S4C**). These data suggest that reduced AQP4 perivascular localization in manifest zQ175 HD mice may mainly be due to relatively lower levels in active astrocytes. Astrogliosis may further exacerbate compromised glymphatic function in HD.

### Reduced AQP4 perivascular localization is evident in human HD brain

The next key question is whether a compromised glymphatic network occurs in the human HD brain. We obtained postmortem caudate putamen samples from NIH Neurobiobank (**Table S1**-fixed tissue samples for immunostaining; **Table S2 –** frozen caudate putamen samples for Western blotting). Our co-immunostaining results with antibodies to AQP4 and collagen IV revealed reduced AQP4 immunosignals in perivascular areas (**Figure 4A**); the quantification of colocalized immunosignals of AQP4 and collagen IV illustrated significantly reduced AQP4 perivascular localization in human HD brains (p<0.05, **Figure 4B**). We then determined protein levels of AQP4, SNTA1, and GFAP in the human HD brain. As we observed in the manifest HD mouse brain, increased astrogliosis indicated by increased GFAP levels was evident in all HD caudate-putamen samples (**Figures 4C, D**). The overall increased AQP4 and SNTA1 signals in Western blots are mainly due to astrogliosis—as when we quantified the ratios of AQP4 and SNTA1 to GFAP, we found that both isoforms of AQP4, M1 and M23, and SNTA1 levels were lower in HD brains than those in control brains (**Figure 4E**). These results demonstrate that the human HD brain presents with an impaired glymphatic network on both the cellular and molecular level.

**Figure 4.**
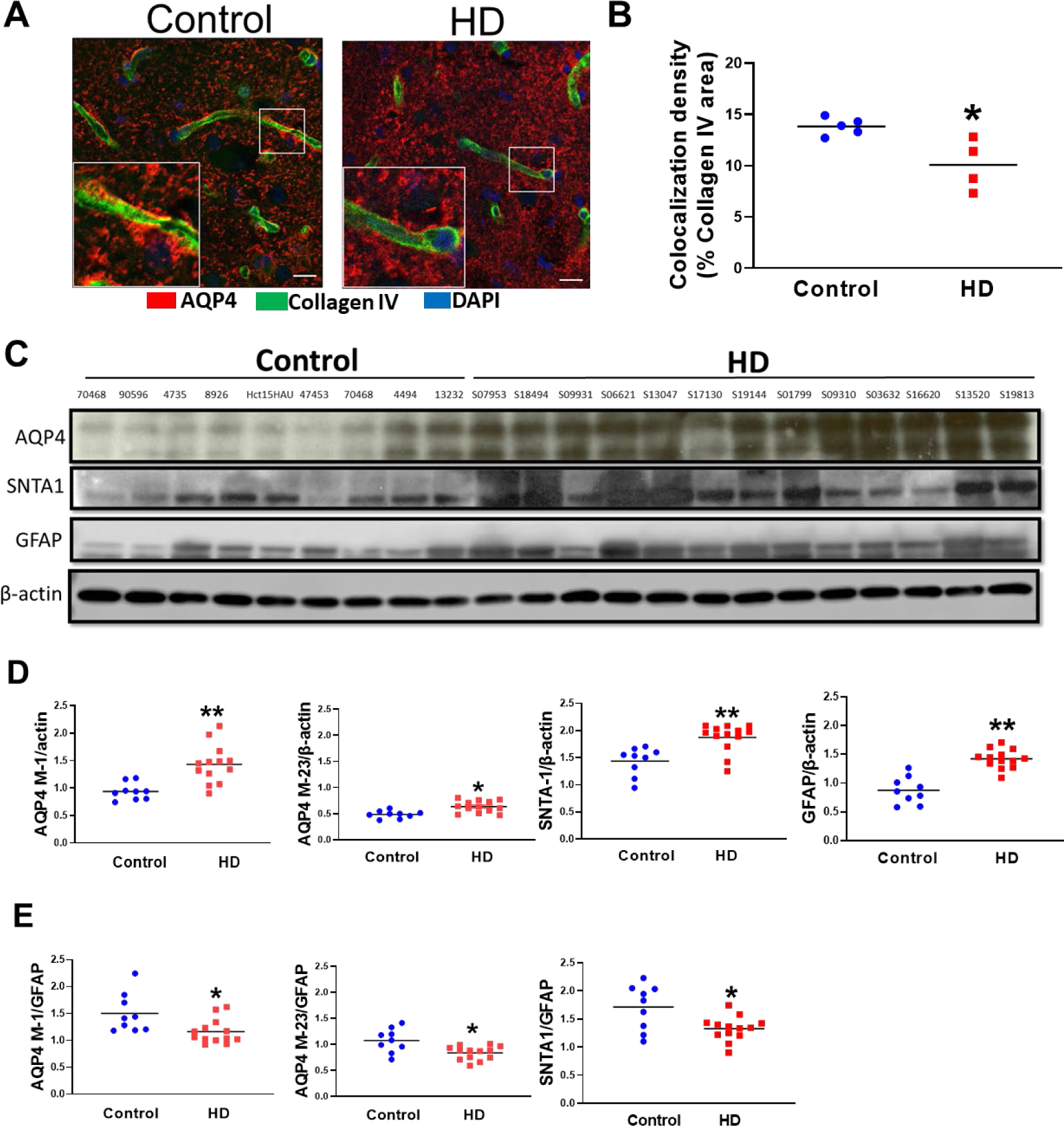
Reduced perivascular AQP4 localization and diminished levels of AQP4 and SNTA1 accompanying astrogliosis in the human HD brain. (**A**) Representative images of co-immunofluorescent staining of AQP4 (red) and collagen IV (green) in the caudate putamen of an HD human subject and age-matched control. (**B**) Quantification of co-localized pixels of AQP4 (red) and collagen IV (green) in the caudate putamen of HD patients (n=6) and age-matched controls (n=4). (**C**) Western blots of AQP4, SNTA1, and GFAP in the human brain from 13 HD brains and 9 control brains. (**D**) Quantification of the ratio of AQP4, SNTA1 to β-actin in the caudate putamen of indicated groups. \*\**p*<0.01 vs. control by Standard Student’s t-test. (**E**) Quantification of ratios of AQP4, SNTA1 to GFAP in the caudate putamen of indicated groups. \**p*<0.05, \*\**p*<0.01 vs. control by Standard Student’s t-test.

## DISCUSSION

In this study, we first assessed glymphatic function with DGE MRI and then validated the findings with gold-standard fluorescent CSF tracer influx measures in a widely used HD knock-in mouse model. Our results demonstrate that zQ175 HD mice exhibited significantly impaired glymphatic function prior to detectable striatal atrophy and motor deficits. A mechanistic experiment suggests that decreased AQP4 levels and its perivascular localization underlie the impaired glymphatic network in premanifest zQ175 HD mice. In both manifest HD mice and postmortem human HD brain, astrogliosis may worsen the already compromised glymphatic clearance capacity.

A fundamental problem in HD and other neurodegenerative disorders is abnormal protein accumulation in the brain, where misfolded and aggregated disease-causative proteins can propagate and spread in a prion-like fashion (*2*). Here we used a recently developed DGE MRI approach, with i.v. D-glucose as a natural biodegradable tracer, to investigate glymphatic function in an HD mouse model. The data show that the glymphatic network is impaired in premanifest HD mice and worsens in manifest HD. Our DGE MRI findings on impairment of glymphatic function in HD mice were further validated by fluorescence-based imaging of CSF tracer distribution in the parenchyma (*3*). Although a recent study suggests that a dysfunctional glymphatic system may interrupt ASO brain distribution (*20*), HD has been far less studied in the context of the glymphatic function. Our present work advances the understanding of a rediscovered brain-wide waste clearance pathway that is functionally interconnected with the brain neurovascular system and its contribution to HD pathogenesis. In the long run, our data provide fundamental insights into the behavior of the glymphatic system in HD and a mechanistic basis for identifying new targets for disease-modifying treatment for HD.

Evidence is building that failure of the glymphatic system leads to an increase of local protein concentrations to levels that favor aggregation. The glymphatic system is a highly organized CSF-ISF transport system with several key functions, including exporting excess metabolite/protein wastes in ISF from brains. The glymphatic system’s perivascular pathways are directly connected to the subarachnoid spaces surrounding the brain, from which CSF is rapidly driven into deep regions of the brain. Regardless of its precise efflux pathways, CSF ultimately drains into the cervical lymphatic vasculature. Caron et al. reported that mHTT is present in the CSF of HD mice in the absence of neurodegeneration (*21*). More specifically, their study showed that secretion of mHTT from cells in the CNS, followed by glymphatic clearance from the extracellular space, contributes to mHTT in the CSF (*21*). The discovery that disease-relevant misfolded and aggregated proteins, such as mHTT, α-synuclein, and Aβ, can propagate and spread in a prion-like fashion has sparked considerable interest (*2*). It has been postulated that seeding occurs across brain regions that are synaptically connected. However, the evidence for synaptic spread is largely based on *post hoc* analysis of anatomic networks; it remains unclear how synaptic relationships by themselves can mediate seeding. It is possible that aggregates simply spread via the extracellular spaces of the glymphatic network, and that aging-and/or disease-relevant gene mutations deteriorate the glymphatic network which could slow the clearance of toxic protein waste. This hypothesis was evident in a mouse AD model, in which deletion of Aqp4 sharply increased both Aβ plaque formation and cognitive deficits (*33*). It might also be the case in HD, though further examination needs to confirm this hypothesis.

Glymphatic functioning is a highly regulated process, with changes in its activity accompanying aging as well as disease conditions (*9-11*). The efficiency of glymphatic clearance is diminished by reduced AQP4 perivascular localization (*12, 13*). There is a growing awareness that reduced perivascular AQP4 occurs in pathological conditions (*14-17*). Deleting AQP4 water channels reduced CSF tracer movement from the periarterial spaces into the interstitium and resulted in markedly impaired clearance of extracellular solutes (*34*). In our study, we detected impaired D-glucose clearance in the CSF of Aqp4 KO mice, and we further observed significantly decreased AQP4 levels and its perivascular localization in the premanifest HD mouse brain which underlies perturbation of glymphatic function. Theories explaining the drainage of brain metabolites via different extracerebral glymphatic pathways are revised now in light of novel findings, including the function of meningeal lymphatics (*6, 7*). Recent data suggest that the glymphatic system and meningeal lymphatic vessels drain macromolecules from the brain parenchyma to the deep cervical lymph nodes (*6, 35*).

The brain is an energy-demanding organ that weighs ∼2% of the whole body but consumes ∼20% of total body D-glucose in the resting state (*36*). Strong evidence of early D-glucose hypometabolism in the HD brain has been reported (*37, 38*). A decreased expression of glucose transporters GLUT1 and GLUT3 was also reported in post-mortem HD brains (*39*). Yet within the brain, when and how much glucose transporters and metabolic changes occur during disease progression remains controversial. D-glucose uptake and metabolism may appear to function differently at distinct disease stages in HD. Whether altered D-glucose uptake and metabolism in premanifest HD represent irreversible damage or compensatory change remains unknown. We found D-glucose uptake was decreased in the manifest zQ175 mouse striatum but not in the CSF, which is consistent with decreased Glut3 levels but not Glut1 levels in the striatum of these HD mice.

The high metabolic rate of the brain requires rapid infusion and clearance of metabolic products because neurons and glial cells are very sensitive to changes in their extracellular homeostasis. The glymphatic system facilitates the convective exchange of various interstitial solutes between CSF and ISF and is important for the brain-wide delivery of nutrients, specifically D-glucose (*40-42*). Altogether, DGE MRI can measure dynamic D-glucose clearance in the CSF, which has the potential to serve as a biomarker of brain glymphatic function. The decline of glymphatic function in premanifest HD condition suggests that treatments directed at restoring normal glymphatic transport and waste removal from the brain may become a preventive or therapeutic approach for this devastating disease.

## MATERIALS AND METHODS

### Animals

All animal experiments were performed in accordance with the Guide for the Care and Use of Laboratory Animals of the National Institute of Health and approved by the Institutional Animal Care and Use Committee (ACUC) at Johns Hopkins University. All mice were housed under standard laboratory conditions and provided with access to food and water. Heterozygous zQ175 mice and wild-type littermates were used. zQ175 breeders were obtained from the Jackson Lab (Bar Harbor, ME).

Genotyping and CAG repeat size was determined by PCR of tail snips at Laragen Inc. (Culver City, CA, USA). The CAG repeat length of zQ175 mice used in the study was 220 ± 3. All mice were housed under specific pathogen-free conditions with a reversed 12-h light/dark cycle maintained at 23°C and provided with food and water ad libitum. All behavioral tests and MRI measures were done in the mouse dark phase (active). During MRI scanning, the mice were under isoflurane anesthesia to minimize suffering (See below for details). Mice were randomized into groups to avoid bias. Data were collected using animal ID and analyzed by investigators who were blinded to genotype and treatment.

### MRI acquisition

In vivo MRI experiments were performed on a horizontal bore 11.7 T Bruker Biospec system (Bruker, Ettlingen, Germany) equipped with a 72 mm quadrature transmitter coil and a four-element (2 × 2) phased-array receiver coil. The MRI detection used an onVDMP sequence [44] consisting of a train of binomial pulses; each pulse shape was two block pulses of opposite phases with a width of 1.5 m s. Peak B1 strength was 15.6 μT. The flip angle of each block pulse was 360°. The offset of the binomial pulses was set to the water resonance, and the mixing time between pulse pairs was set to 10ms to remove unwanted traverse magnetization. After the labeling module, anatomical images covering the third ventricle were acquired using a multi-slice Rapid Acquisition with Relaxation Enhancement (RARE) sequence: resolution=0.1× 0.1 × 0.5 mm3, echo time (TE)=3ms, repetition time (TR)=2.5s, mixing time (TM)=10ms, average number (NA)=128, RARE factor = 23. Slice thickness = 1 mm, a matrix size of 196 × 128, and a field-of-view (FOV) of 20 × 10 mm2. A multi-slice approach was used to observe glucose uptake in both the striatum and CSF. DGE images were acquired continuously for 50 min. A 5min pre-scan was performed before glucose infusion as dummy scans to ensure the mouse was stabilized in the scanner, then a bolus of 0.15 mL 50% w/w D-glucose (0.5 g/mL, clinical-grade dextrose, Hospira, Lake Forest, IL) was infused (rate, 0.15 ml/min) through the tail vein using a syringe pump (Harvard Apparatus, Holliston, MA, USA). The amount of glucose for each mouse depended on body weight (6μl/g).

All mice were anesthetized using 2% isoflurane and maintained by 1-1.5% isoflurane during the MRI scan. The mouse was placed on a water-heated bed equipped with a pressure sensor. The animal head was positioned with a bite bar and a container. The respiratory rate of the mouse was monitored (SAII, Stony Brook, NY, USA) and maintained at 40–60 breaths/min.

### DGE MRI with onVDMP sequence image analysis

Statistical parametric mapping (SPM) (Version 8, Wellcome Trust Centre for Neuroimaging, London, United Kingdom; http://www.fil.ion.ucl.ac.uk/spm/) and in-house programs coded in Matlab (MathWorks, Natick, MA, USA) were used. DGE MRI was measured with the onVDMP pulse sequence consisting of a train of binomial pulses composed of two high-power pulses with alternating phase (*pp*) (*22*). The binomial pulse is designed to label all exchanging protons in a way that depends on the excitation profile, designed to avoid water resonance. Motion correction between DGE images was performed with Medical Imaging Registration Toolbox, and the difference in signal was quantified with the surround subtraction method. CEST approaches such as onVDMP MRI can also be described using T1ρ theory. The baseline signal and curve fitting follow our previous publications^26^, ^41^ on DGE MRI. Two regions of interest (ROIs) were manually selected in each scan: 1) CSF (the third ventricle) and 2) striatum. The measurement of time-resolved changes of DGE MRI signal after an intravenous bolus infusion of D-glucose reports changes in glucose concentrations in biological tissues and thus contains information on D-glucose delivery, transport, metabolism, as well as clearance kinetics.. The measurement of time-resolved changes of DGE MRI signal after an intravenous bolus infusion of D-glucose reports changes in glucose concentrations in biological tissues and thus contains information on D-glucose delivery, transport, metabolism, as well as clearance kinetics.

This molecular MRI method allows high labeling efficiency by utilizing a train of short pulses leading to an improved Chemical Exchange Saturation Transfer (CEST) signal for glucose hydroxyl protons compared to conventional CEST MRI. To facilitate the translation of this DGE MRI to clinical use, we have tested the onVDMP pulse sequence to detect D-glucose CSF clearance on a 3T scanner, a common clinical MRI field strength (*26*). We also showed the feasibility of this modification in other neurodegenerative disorders, including two mouse models of AD (*26, 43*). Glucose transporters are highly enriched in brain capillary endothelial cells; thus, D-glucose can easily penetrate the BCSFB and enter CSF. This provides an opportunity to monitor the CSF-ISF exchange process through the intravenous administration of D-glucose, a minimally invasive method that is a routine clinical practice.

### Intracisternal tracer injection

Anesthetized mice with the same concentration of isoflurane used in MRI were fixed in a stereotaxic frame, the cisterna magna (CM) was surgically exposed, and a 30GA needle was inserted into the cisterna magna. The fluorescent CSF tracer (bovine serum albumin, Alexa Fluor 647 conjugate, Invitrogen, Life Technologies, Eugene, OR, USA; 66 kDa) was dissolved in artificial CSF at a concentration of 0.5% (w/v). 10µl of CSF tracer was injected at a rate of 2µl/min for 5 minutes using a syringe pump (Harvard Apparatus). All experiments were conducted by the same operator. To visualize tracer movement from the subarachnoid space into the brain parenchyma, mice were perfusion fixed 1 hour after intracisternal tracer injection. 100µm coronal vibratome slices were cut and mounted as above, and tracer influx into the brain was imaged ex vivo by a fluorescence microscope (Zeiss), with the tile function to rebuild the whole slice. Tracer influx figures were quantified independently by two sets of blinded investigators using Fiji (Image J) software, as described in a previous study. Each coronal slice was manually outlined, and the mean fluorescence intensity was measured at a constant threshold. Average fluorescence intensity was calculated across 4 slices for a single animal. Equivalent slices were used for all biological replicates.

### Immunohistochemistry

Mice were anesthetized with isoflurane and perfused transcardially with phosphate-buffered saline (PBS) followed by 4% paraformaldehyde. Brains were post-fixed overnight, followed by immersion in 30% sucrose for 24 h. Coronal sections (40 μm) were cut on a microtome. Sections were immunostained with primary antibodies, AQP4 (249323, 1:100, Alomone Labs, Israel), and Collagen Ⅳ (2150-1470, 1:100, BioRad, USA). Briefly, the sections were washed three times with PBS (10 min each time), then permeabilized by incubating with 0.3% Triton X-100 for 5 min, followed by incubation with blocking solution containing 3% donkey serum, 3% goat serum, and 0.3% Triton X-100 for 1 h. The sections were then incubated with primary antibody at 4°C overnight. After three washes with PBS, the sections were incubated with fluorescence-labeled secondary antibody for 2 h at room temperature and then washed 3 times with PBS. Sections were mounted onto Superfrost slides (Fisher Scientific, Pittsburgh, PA, USA), dried, and then covered with anti-fade mounting solution. Fluorescence images were acquired with an LSM 700 AxioObserver fluorescence microscope.

For image analysis, the samples were coded with ID, and the images were analyzed by investigators blinded to genotypes and patient information. Results were then calculated statistically and decoded by different investigators at the end. The results from three microscopy fields per slide and three sections per mouse were calculated for each mouse brain. The colocalization quantities (AQP4+ Collagen Ⅳ+) were determined using the analyze particles plugin function of ZEN3.4 (blue edition). The numbers of AQP4 and Collagen Ⅳ positive puncta in the field (160×160 µm2) were determined at a constant threshold for each stain using × 20 images for quantifications.

### Western Blotting

Striatal tissue samples were homogenized in a buffer containing 50 mM Tris-HCl, pH 8.0, 150 mM NaCl, 0.1% (w/v) SDS, 1.0% NP-40, 0.5% sodium deoxycholate, and 1% (v/v) protease inhibitors mixture. For SDS PAGE, 20 μg of proteins were separated in a 4–20% gradient gel and transferred to a nitrocellulose membrane. The membrane was blotted with the following primary antibodies: AQP4 (249323, 1:100, Alomone Labs, Israel), SNTA-1 (PA5-77702, 1:1000, Thermo Fisher, USA), GFAP (13-0300, 1:1000, Thermo Fisher, USA), GLUT1 (SPM498, 1:1000, Thermo Fisher, USA), Glut3(MA5-32697, 1:1000, Thermo Fisher, USA) and mouse anti-β-actin (Sigma, mouse monoclonal antibody, 1:5000). After incubation with HRP-conjugated secondary antibodies, bound antibodies were visualized by chemiluminescence. The intensity of the Western blot bands was quantified by Image J software.

### Statistics

Data are expressed as the mean ± SEM unless otherwise noted. Student’s t-test was used to measure the significant levels between WT or Control and zQ175/+ or human HD groups at each given timepoint. P-values less than 0.05 were considered statistically significant. Technical replication is represented by n in the figure legends.

## Data and materials availability

All data associated with this study are present in the paper or the Supplementary Materials. Additional requests for raw and analyzed data or materials should be made to and will be promptly reviewed to determine whether the application is subject to any intellectual property or confidentiality requirements. Any data and materials that can be shared will be released after the execution of a material transfer agreement.

## Acknowledgement

We thank NIH NeuroBioBank for providing human samples for this study.

## Author contributions

Conceptualization W.D., H.L., J.X.; Methodology W.D., H.L., C.L., J.X.; Experimental Work H.L., L.C., C.Z., C.L., Y.L., L.C., Z.W., Z.Z.; Writing-original Draft. W.D., H.L., J.X.; Writing-Review & Editing, W.D., H.L., C.L., J.X., H.L., P.C.M.Z., J.J.I; Funding Acquisition, W.D. and J.X.; Supervision, W.D. and J.X.

## Supplementary Materials

## Materials and Methods

### Animals

Aqp4 KO mouse breeders were kindly provided by Dr. Quigley’s lab at Johns Hopkins; the original line of Aqp4 KO mice was generated by Dr. Ole Petter Ottersen using the Genoways technique as described previously (Thrane et al., 2011). Their strategy involved cloning and sequencing a targeted region of the murine Aqp4 gene in a 129/Sv genetic background. Identification of a targeted locus of the Aqp4 gene permitted the deletion of exons 1–3 to avoid any expression of putative splice variants. Hence, a flippase recognition target (FRT)-neomycin-FRT-LoxP–validated cassette was inserted downstream of exon 3 and a LoxP site was inserted upstream of exon 1 (Thrane et al., 2011). The mice were backcrossed for 20 + generations with C57BL/6N mice prior to experimentation.

### *In Vivo* structural MRI acquisition

*In vivo* MRI was performed on a vertical bore 9.4 Tesla MR scanner (Bruker Biospin, Billerica, MA, USA) with a triple-axis gradient and a physiological monitoring system (EKG, respiration, and body temperature). Mice were anesthetized with isoflurane (1%) mixed with oxygen and air at 1:3 ratios via a vaporizer and facial mask and scanned longitudinally (the same mice were imaged repeatedly over 12 months). We used a 20-mm diameter volume coil as the radiofrequency transmitter and receiver. The temperature was maintained by a heating block built into the gradient system. Respiration was monitored throughout the entire scan.

High-resolution anatomical images were acquired using a three-dimensional (3D) T2-weighted fast spin echo sequence with the following parameters: echo time (TE)/repetition time (TR) = 40/700 ms, resolution = 0.1 mm × 0.1 mm × 0.25 mm, echo train length = 4, number of averages = 2, and flip angle = 40°. Multi-slice T2-weighted images of the mouse brain were acquired by the RARE (Rapid Acquisition with Refocused Echoes) sequence with the following parameters (echo time (TE) / repetition time (TR) = 40 ms/1500 ms, RARE factor = 8, in-plane resolution = 0.125 mm x 0.125 mm, slice thickness = 1 mm, total imaging time less than 2 min) and used for high-resolution anatomical imaging. The total imaging time was about 50 min per mouse. Mice recovered quickly once the anesthesia was turned off, and all mice survived the imaging sessions.

### Structural MRI image analysis

Images were first rigidly aligned to a template image using automated image registration software (http://bishopw.loni.ucla.edu/AIR5/, AIR). The template image was selected from one of the images acquired from age-matched littermate control mice (the mouse had the medium brain volume among the control group), which had been manually adjusted to the orientation defined by the Paxinos atlas with an isotropic resolution of 0.1 mm x 0.1 mm x 0.1 mm per pixel. After rigid alignment, images had the same position and orientation as the template image, and image resolution was also adjusted to an isotropic resolution of 0.1 mm × 0.1 mm × 0.1 mm per pixel. Signals from non-brain tissue were removed manually (skull-stripping). Skull-stripped, rigidly aligned images were analyzed by using Landmarker software (www.mristudio.org). Intensity values of the gray matter, white matter, and cerebral spinal fluid were normalized to the values in the template images using a piece-wise linear function. This procedure ensured that the subject image and template image had similar intensity histograms. The intensity-normalized images were submitted by Landmarker software to a Linux cluster, which runs Large Deformation Diffeomorphic Metric Mapping (LDDMM). The transformations were then used for quantitative measurement of changes in local tissue volume among different mouse brains by computing the Jacobian values of the transformations generated by LDDMM (Zhang et al., 2010). There are 29 different brain regions segmented automatically.

### D-Glucose uptake image analysis

The average water signal intensity before infusion is S_base_. Motion correction was conducted using Medical Imaging Registration Toolbox (Myronenko et al., 2010). The regions of interest (ROIs) were manually selected according to the mouse brain atlas (https://mouse.brain-map.org). The glucose uptake curves detected by ^1^H MRS and DGE MRI were fitted by exponential functions to make a quantitative comparison (Huang et al):

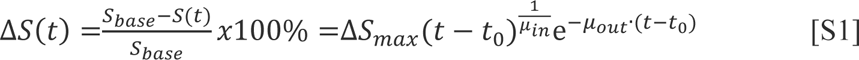

Where ΔS(t) is the signal difference determined by DGE MRI. μ_in_ and μ_out_ are the glucose uptake and outflow rates, respectively. ΔS_max_ represents the maximum signal difference for DGE. A two-sample *t*-test was conducted on the fitted parameters between the HD and WT groups.

### Colocalization density analysis

ZEN 3.4(Black edition) was used to do the Colocalization analysis. The threshold of CH1 is set to 80 to define the green signal (collagen IV). The threshold of CH2 is set to 120 to define the red signal (AQP4). In this way, the Collagen IV area (green pixels), the AQP4 area (red pixels) and the colocalized area(yellow pixels) are generated. Colocalization density was quantified as the percentage of collagen IV and AQP4 positive immunofluorescence within the collagen IV area (yellow pixels / green pixels).

### Behavioral tests

5mm balance beam testing was conducted on an 80 cm long and 5 mm wide square-shaped balance beam that was mounted on supports of 50 cm in height. A bright light illuminated the start platform, and a darkened enclosed 1728 cm^3^ escape box (12 × 12 × 12 cm^3^) was situated at the end of the beam. Mice were trained to walk across the beam twice at least 1 h prior to testing. The time for each mouse to traverse the balance beam was recorded with a 125-sec maximum cut-off, and falls were scored as 125 sec.

**Figure S1.**
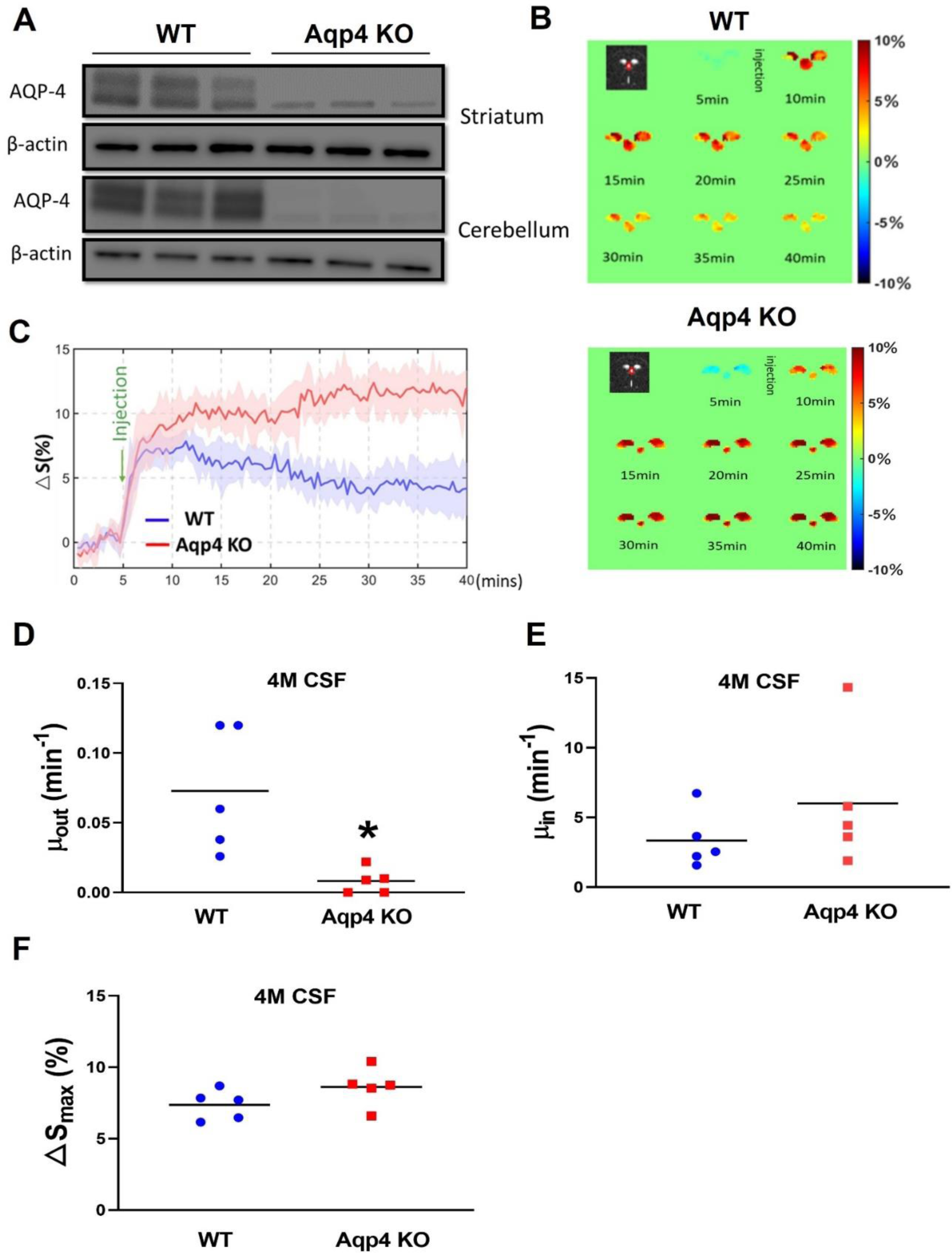
DGE MRI detects perturbed glymphatic function in Aqp4 KO mice at 4 months. (**A**) Western blots of AQP4 in the striatum and cerebellum of Aqp4 KO mice and wild-type (WT) controls. (**B**) Representative DGE images (signal described in Eq. S1) as a function of time in a WT mouse (upper panel) and an Aqp4 KO mouse (lower panel) at 4 months of age. (**C**) The average dynamic D-glucose signals in CSF during the entire scan period from WT (n=5) and Aqp4 KO (n=5) mice. (**D-F**) Comparison of fitted uptake parameter μ_in_ (D), clearance parameter μ_out_ (E) and maximal glucose levels ΔS_max_ (F) between WT and Aqp4 KO mice. \**p*<0.05 vs. WT by Standard Student’s t-test.

**Figure S2.**
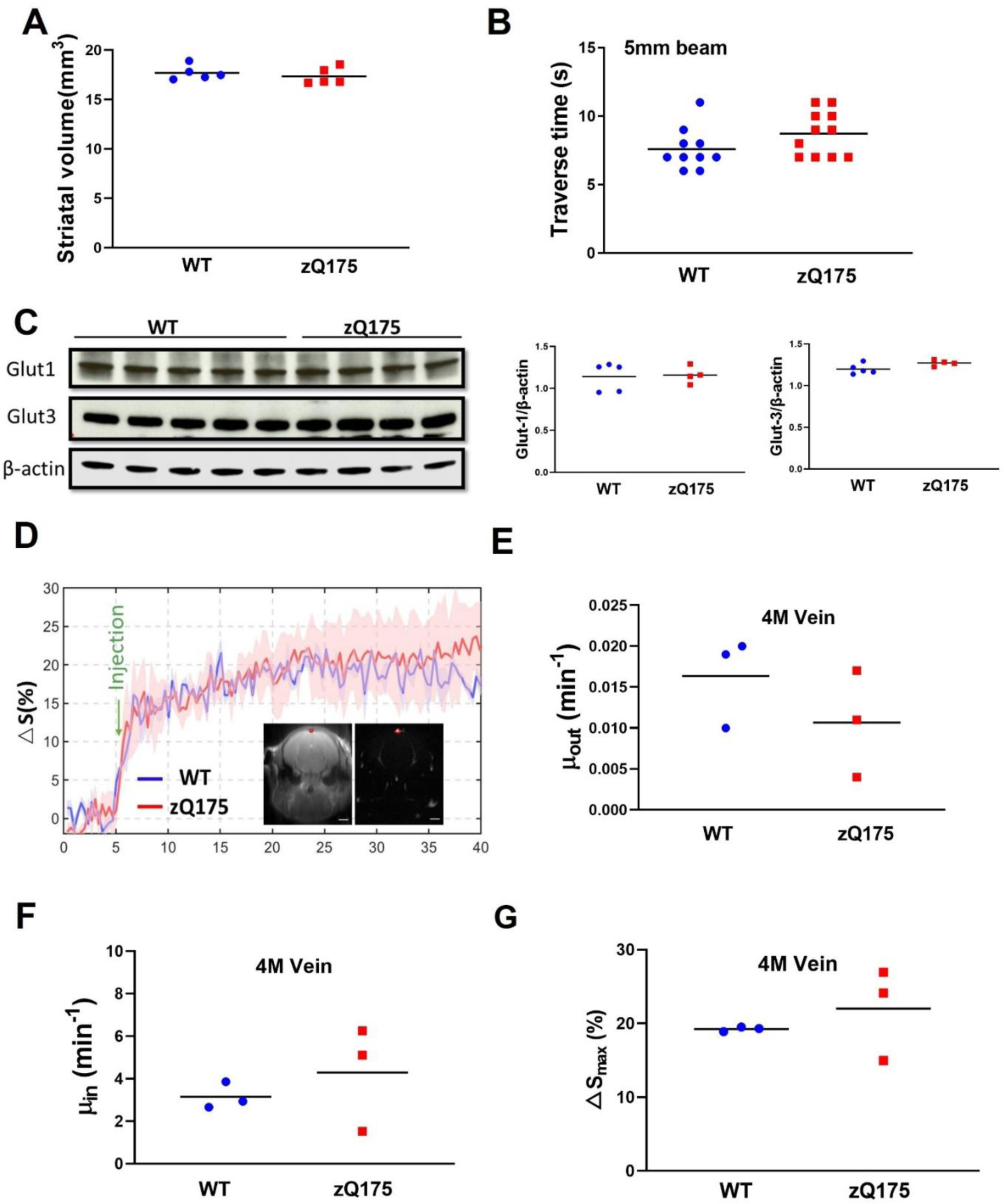
No difference in the striatal volume, motor function, and D-glucose uptake and clearance in the venous blood (sagittal sinus) between 4-month-old zQ175 and WT mice. (**A**) Striatal volume measured by structural MRI in zQ175 and age-matched WT mice. n=5 mice/group. (**B**) Motor coordination assessment on a 5mm balance beam, the traverse time was recorded from WT (n=9) and zQ175 (n=11) mice. (**C**) Western blots and quantification of glucose transporters Glut1 and Glut3 in the striatum of mice. (**D**) DGE curves for the sagittal sinus (red ROI) in WT and zQ175 mice. ROI within the blood vessel (red circle). (**E-G**) D-glucose clearance rate (E), uptake rate (F), and maximal glucose levels in the sagittal sinus vein following i.v injection of D-glucose in the tail vein (G).

**Figure S3.**
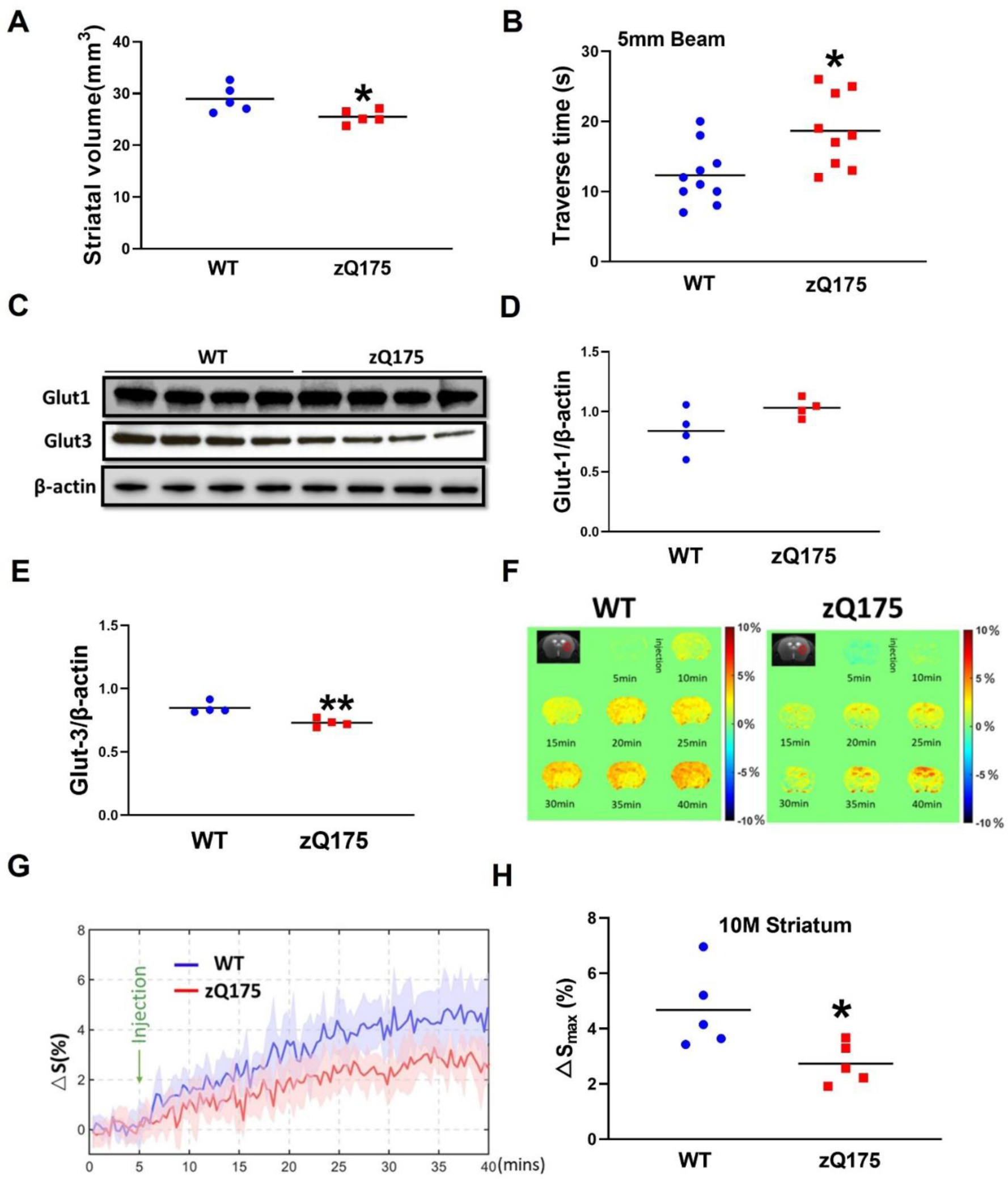
Reduced striatal volume, motor deficits, and perturbed D-glucose uptake in the striatum are evident in 10-month-old zQ175 mice. (**A**) Striatal volume measured by structural MRI in zQ175 and age-matched WT mice. n=5 mice/group. \**p*<0.05 vs WT by Standard Student’s t-test. (**B**) Motor coordination assessment on a 5mm balance beam, the traverse time was recorded from WT and zQ175 mice. n=10, \**p*<0.05 vs. WT by Standard Student’s t-test. (**C**) Western blots of Glut1 and Glut3 in the striatum of WT and zQ175 mice. (**D-E**) Quantification of Glut1 and Glut3 levels from Western blots in C. n=4, \*\**p*<0.01 vs. WT by Standard Student’s t-test. (**F**) Representative images of D-glucose uptake in the striatum of WT and zQ175 mice. The hot color indicates higher uptake. (G) The DGE curves of D-glucose uptake following i.v. injection in the striatum of WT and zQ175 mice. n=5. Note decreased D-glucose uptake in HD mouse striatum. (**H**) The maximal D-glucose concentrations in WT and HD mouse striatum after i.v. infusion. n=5, \**p*<0.05 vs. WT by Standard Student’s t-test.

**Figure S4.**
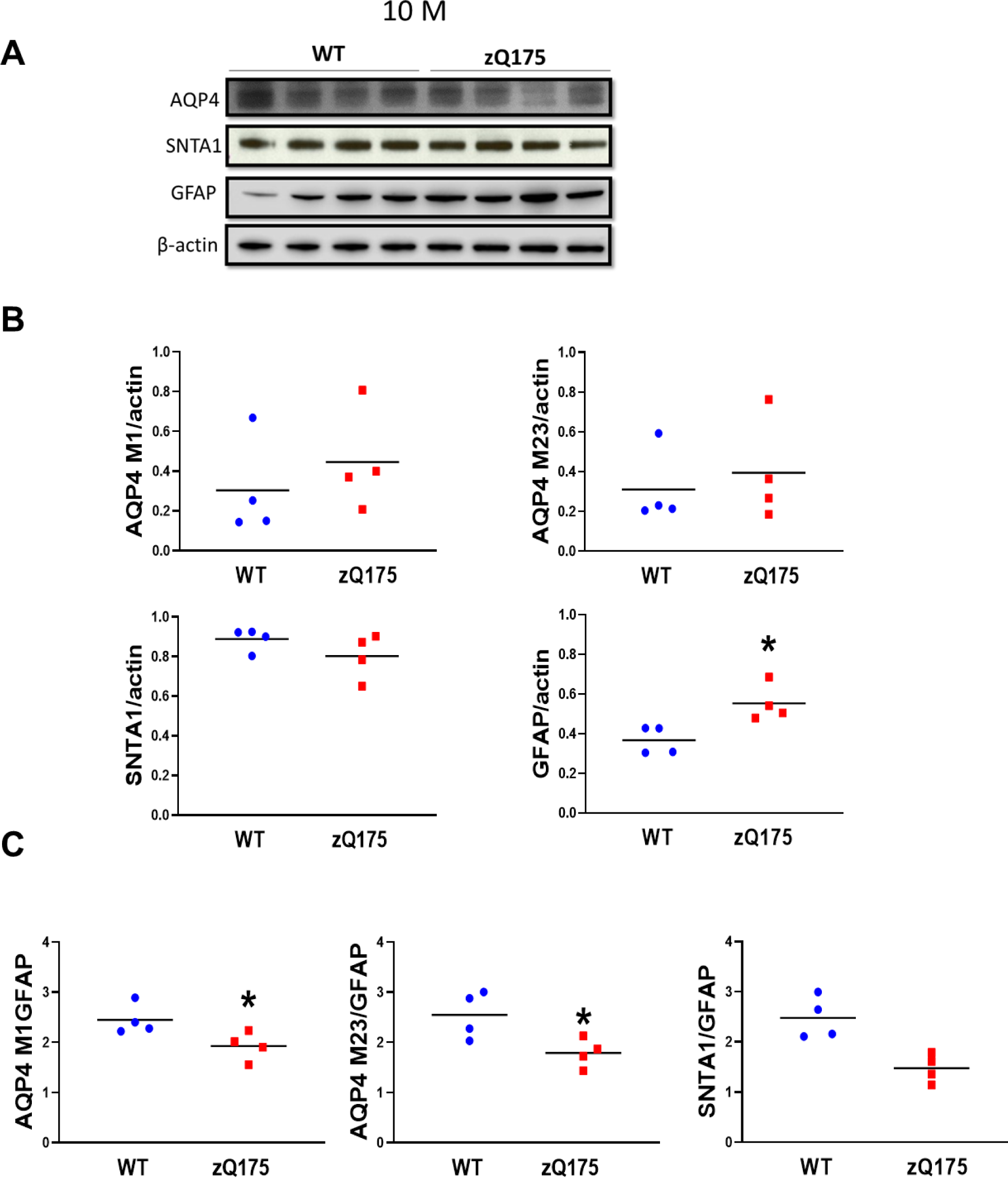
Progressive astrogliosis exacerbates the disturbed molecular network of the glymphatic system in manifest zQ175 HD mice. **(A)** Western blots images of samples from 10-month-old mice. (**B**) Quantification of AQP4, SNTA1, GFAP in the striatum of 10-month-old zQ175 mice and WT controls. \**p*<0.05 vs. WT by Standard Student’s t-test. (**C**) Ratios of AQP4 M1, AQP4 M23 and SNTA1 to GFAP in the striatum of 10-month-old zQ175 mice and WT controls. \**p*<0.05 vs. WT by Standard Student’s t-test.

**Table S1.** Postmortem caudate putamen samples from NIH Neurobiobank for immunostaining.

|  | Subject ID | Age | Brain Region | Subject Sex | Clinical Brain Diagnosis | Preparation |
| --- | --- | --- | --- | --- | --- | --- |
|  | S04230 | 80 | Caudate nucleus | Male | Huntington's disease | 4% Paraformaldehyde Fixed |
|  | S08214 | 93 | Caudate nucleus | Male | Huntington's disease | 4% Paraformaldehyde Fixed |
|  | S08434 | 70 | Caudate nucleus | Male | Unaffected Control | 4% Paraformaldehyde Fixed |
|  | S09101 | 72 | Caudate nucleus | Male | Huntington's disease | 4% Paraformaldehyde Fixed |
|  | S13047 | 72 | Caudate nucleus | Male | Huntington's disease | 4% Paraformaldehyde Fixed |
|  | S16578 | 71 | Caudate nucleus | Male | Unaffected Control | 4% Paraformaldehyde Fixed |
|  | S16640 | 70 | Caudate nucleus | Male | Unaffected Control | 4% Paraformaldehyde Fixed |
|  | S16891 | 71 | Caudate nucleus | Male | Unaffected Control | 4% Paraformaldehyde Fixed |
|  | S18965 | 69 | Caudate nucleus | Male | Unaffected Control | 4% Paraformaldehyde Fixed |

**Table S2.** Postmortem caudate putamen samples from NIH Neurobiobank for Western blotting.

|  | Subject ID | Age | Brain Region | Subject Sex | Clinical Brain Diagnosis | Preparation |
| --- | --- | --- | --- | --- | --- | --- |
|  | 70468 | 64 | Caudate nucleus | Male | Unaffected Control | Frozen |
|  | 90596 | 66 | Caudate nucleus | Male | Unaffected Control | Frozen |
|  | 4735 | 73 | Caudate nucleus | Male | Unaffected Control | Frozen |
|  | 8926 | 65 | Caudate nucleus | Male | Unaffected Control | Frozen |
|  | Hct15HAU | 65 | Caudate nucleus | Male | Unaffected Control | Frozen |
|  | 47453 | 55 | Caudate nucleus | Male | Unaffected Control | Frozen |
|  | 70468 | 64 | Caudate nucleus | Male | Unaffected Control | Frozen |
|  | 4494 | 67 | Caudate nucleus | Male | Unaffected Control | Frozen |
|  | 13232 | 65 | Caudate nucleus | Male | Unaffected Control | Frozen |
|  | S07953 | 71 | Caudate nucleus | Male | Huntington Grade 2 | Frozen |
|  | S18494 | 71 | Caudate nucleus | Male | Huntington Grade 2 | Frozen |
|  | S09931 | 75 | Caudate nucleus | Male | Huntington Grade 2 | Frozen |
|  | S06621 | 75 | Caudate nucleus | Male | Huntington Grade 2 | Frozen |
|  | S13047 | 72 | Caudate nucleus | Male | Huntington Grade 2 | Frozen |
|  | S17130 | 76 | Caudate nucleus | Male | Huntington Grade 2 | Frozen |
|  | S19144 | 73 | Caudate nucleus | Male | Huntington Grade 3 | Frozen |
|  | S01799 | 71 | Caudate nucleus | Male | Huntington Grade 3 | Frozen |
|  | S09310 | 70 | Caudate nucleus | Male | Huntington Grade 3 | Frozen |
|  | S03632 | 75 | Caudate nucleus | Male | Huntington Grade 3 | Frozen |
|  | S16620 | 71 | Caudate nucleus | Male | Huntington Grade 3 | Frozen |
|  | S13520 | 71 | Caudate nucleus | Male | Huntington Grade 3 | Frozen |
|  | S19813 | 70 | Caudate nucleus | Male | Huntington Grade 3 | Frozen |

